# Temporal phosphoproteomics reveals rapid restoration of kinase signaling by *Glycyrrhiza glabra* in a rotenone-induced Parkinson disease model

**DOI:** 10.64898/2026.06.15.732239

**Authors:** Vanya Kadla Narayana, Gayathree Karthikkeyan, Mohd. Altaf Najar, Ravishankar Pervaje, Thottethodi Subrahmanya Keshava Prasad, Prashant Kumar Modi

**Affiliations:** Center for Systems Biology and Molecular Medicine, Yenepoya Research Centre, Yenepoya (Deemed to be University), Mangalore 575018, India; Sushrutha Ayurveda Hospital, Puttur 574201, India; Center for Omics & Systems Medicine (C-OSM), Nitte (Deemed to be University), Mangalore 575018, India

**Keywords:** Ayurveda, Traditional Medicine, Actionable molecules, Signaling pathways

## Abstract

Parkinsons disease is a progressive neurodegenerative disorder associated with mitochondrial dysfunction, oxidative stress, impaired autophagy, and dysregulated cellular signaling pathways. Although *Glycyrrhiza glabra* has been reported to exhibit neuroprotective properties, the early phosphorylation-mediated signaling mechanisms underlying its protective effects remain poorly understood. In this study, we employed a Tandem Mass Tag (TMT)-based temporal quantitative phosphoproteomic approach to investigate early signaling events associated with *Glycyrrhiza glabra*-mediated neuroprotection in a rotenone-induced in vitro PD model. Differentiated IMR-32 neuronal cells were treated with rotenone alone or in combination with *Glycyrrhiza glabra* extract, and phosphoproteomic alterations were analyzed at 2, 5, 15, and 30 minutes using liquid chromatography coupled with tandem mass spectrometer. Temporal phosphoproteomic analysis identified 6,424 phosphopeptides corresponding to 2,368 phosphoproteins and 5,468 phosphorylation sites. Comparative analysis revealed extensive phosphorylation rewiring induced by rotenone and restoration of several dysregulated phosphorylation events following *Glycyrrhiza glabra* co-treatment. More than 130 phosphoproteins and multiple kinase-associated signaling pathways were dynamically regulated across the temporal conditions. Kinase enrichment analysis identified restoration of several critical kinases, including *AKT1, MTOR, MAPK1/3, PRKACA, PRKCD, and GSK3A/B*, which are associated with neuronal survival, stress adaptation, and autophagy. Integrated pathway and kinase-substrate interaction analyses further revealed enrichment of AMPK signaling, FOXO signaling, receptor tyrosine kinase signaling, RNA processing, and cell-cycle regulatory pathways. Notably, several spliceosome-associated phosphoproteins demonstrated dynamic phosphorylation changes during the early neuroprotective response. Collectively, this study provides a detailed temporal phosphoproteomic landscape of early signaling events associated with *Glycyrrhiza glabra*-mediated neuroprotection and highlights kinase-driven signaling pathways that may represent potential therapeutic targets in Parkinsons disease.

## Introduction

Parkinson’s Disease (PD) is a progressive neurodegenerative disordermarked by the death of dopaminergic neurons in the substantia nigra pars compacta (SNpc) and characterized by tremor, rigidity, and late-stage postural instability (**1**). The prevalence of PD will be estimated at 267 cases per 100,000 in 2050 (**2**). The hallmarks of PD include endoplasmic reticulum (ER) stress, accumulation of misfolded alpha-synuclein (aSyn), PINK1/parkin, receptor-mediated mitophagy leading to neuronal death, and reduced dopamine release into the striatum, resulting in motor loss (**3–8**).. The management of PD involves the use of levodopa, a commonly prescribed medicine, which is helpful in the disease management in early stages; however, continued use of this medicine is often reported to result in decreased efficacy of the medicine and is often associated with many side effects, leading to compromising the quality of life of the patients (**9,10**).. Given the complex nature of PD, current treatment relies on symptomatic relief and does not cure the disease. This underscores the growing need for complementary therapeutic approaches that target early pathogenic events in PD. Therefore, a sustainable alternative and complementary management strategy for PD is necessary, which not only mitigates PD symptoms but also rectifies the underlying molecular aberrations and enhances quality of life.

The search for long-term disease management prompted an investigation into complementary and alternative medicines, which integrate traditional practices. As an alternative strategy for the management of PD, the traditional Complementary and Alternative Medicine (CAM) is explored (**11**). One such system, the Indian Ayurvedic medicinal system, categorizes medicinal plants having nootropic characteristics as Medhya Rasayana, which enhances memory and cognitive abilities (**12**). Medhya Rasayana includes Yashtimadhu (*Glycyrrhiza glabra*, common name: licorice), Mandukaparni (*Centella asiatica*, common name: Asiatic pennywort), Guduchi (*Tinospora cordifolia* (Wild) Miers), Shankhapushpi (*Convolvulus pleuricaulis* Chois), and Brahmi (*Bacopa monnieri*, common name: water hyssop) (**12–15**).

*Glycyrrhiza glabra* (*G. glabra*, Yashtimadhu) powder is prepared from the dried roots of *Glycyrrhiza glabra* L, generally referred to as licorice (http://www.theplantlist.org/; http://www.ayurveda.hu/api/API-Vol-1.pdf). It is classified as a Medhya Rasayana, with memory-enhancing, antidepressant, and antioxidant properties (**14,16–21**). We have previously shown the neuroprotective activity of *G. glabra* powder in a rotenone-induced PD model using mass spectrometry (MS)-based proteomic and metabolomic analyses (**16,22**). However, the role of *G. glabra-*mediated regulation of phosphorylation dynamics in phosphoproteins, and its effects on kinases in PD, remain underexplored. This study sheds light on the kinase-substrate signaling network that regulates the early pathogenic phosphorylation events occurring in rotenone-treated cells that are restored by *G. glabra* co-treatment.

Several studies have identified the kinase signaling network associated with PD pathogenesis and emphasized the need to develop new treatments for the management of PD (**23–29**). The dysregulation of kinases such as Mitogen-Activated Protein Kinases, Phosphatidylinositol 3-Kinase (PI3K)/ Protein Kinase B (AKT), Mechanistic Target of Rapamycin Kinase (MTOR), Janus Kinases (JAK) and Glycogen Synthase Kinases (GSK) results in PD pathogenesis (**29–33**). The tight regulation of these kinases is essential for the proper functioning of the neurons and critical forneuronal survival. The kinase activity is regulated by their phosphorylation states, which can be effectively assessed using MS-based phosphoproteomic analysis. MS-based phosphoproteomic analysis employs the enrichment and study of the phosphopeptides using affinity-based methods. Understanding the dynamics of protein phosphorylation highlights the kinases-substrates links involved in signaling events and their downstream effects.

Several phosphoproteomic studies have previously demonstrated the signaling kinases and their networks involved in the PD pathogenesis (**34–38**)and the neuroprotection conferred by small molecules (**39,40**).

Despite the availability of this powerful technology, its application in elucidating the neuroprotective targets of traditional and alternative medicines remains limited. Therefore, in this study, we sought to explore the mechanistic and temporal events involved in the neuroprotection mediated by *G. glabra* against a rotenone-induced in vitro model of PD, using temporal quantitative total proteomic and phosphoproteomic approaches. We describe the temporal signaling events and important kinase-substrate associations that may serve as novel *G. glabra* targets involved in its neuroprotective mechanism.

## Materials and methods

### Materials procurement

Rotenone (Cat. No. R8875), retinoic acid (Cat. No. R2625), collagen (Cat. No. C9791), iodoacetamide (Cat. No. I6125), and DL-dithiothreitol (Cat. No. D9779) were procured from Sigma-Aldrich, St. Louis, USA. Dulbecco’s Modified Eagle Medium-high glucose (DMEM, Cat. No. 12100046), fetal bovine serum (FBS, Cat No 10270-106), Trypsin EDTA (Cat. No. 25200-056), and 100X antibiotic/antimycotic solution (Cat. No. 15240062) were purchased from Gibco, Thermo Fisher Scientific USA. Pierce™ BCA protein estimation assay kit (Cat. No. 23225), Pierce™ Peptide estimation assay kit (Cat. No. 23275), and TMT 10plex kit (Cat. No. 90110), High-Select TiO2 (Cat. No. A32993) and High-Select Fe-NTA (Cat. No. A32993) Phosphopeptide enrichment kits, were procured from ThermoFisher Scientific USA. TPCK-treated Trypsin (Cat. No. LS003741), from Worthington Biochemical Corporation, USA. Sep-Pak C-18 (Cat. No. WAT051910) was procured from Waters, USA.

### *G. glabra* procurement and preparation of *G. glabra* extract

*Glycyrrhiza glabra* L. is commonly known as licorice/liquorice and in India as Yashtimadhu (http://www.theplantlist.org/).*G. glabra* powder (lot no. 64) was procured from a GMP-certified manufacturer of Ayurvedic formulations, SDP Remedies and Research Centre, Puttur, Karnataka, India. The plant was authenticated by Dr. Harikrishna Panaje, SDP Remedies and Research Centre microscopely and macroscopically. A specimen of *G. glabra* roots is maintained in the centre (ID: SDP/YM/001-2017). The dry roots of *G. glabra* are used for the preparation of the powder. The powder was prepared as previously described (**16,41**). The *G. glabra* root powder was authenticated using LC-MS/MS analysis for the presence of the lead bioactive molecules, including glycyrrhizic acid, glabridin (specific to *G. glabra*), licoricesaponin-G2, and liquiritin apioside, in our previous study (**41**). *G. glabra* aqueous extract for treatment was prepared as described (**16**). Briefly, *G. glabra* powder was suspended in Milli-Q water at a concentration of 0.1 g/ml, in a final volume of 10 ml. The aqueous extraction mixture was incubated overnight at room temperature under continuous rotation. The extract was centrifuged at 5,000 rpm for 10 minutes, and the supernatant was collected. The aqueous extract was subsequently dried using Speed Vac (Savant, Thermo Fisher Scientific, USA) and stored at -20 °C until further use. For cell culture experiments, the dried extract was reconstituted in serum-free culture media.

### Cell culture treatment

IMR-32 cells (ATCC® CCL-127™) were procured from the National Centre for Cell Sciences, Pune, India. Changes in protein phosphorylation dynamics were assessed at early time points of 2, 5, 15, and 30 minutes. The IMR-32 cells were seeded at a density of 1.5 × 10^5^ cells/plate onto 10-cm culture plates and maintained in Dulbecco’s Modified Eagle Medium (DMEM)-high glucose media, supplemented with 10% fetal bovine serum (FBS), and 1X antibiotic/antimycotic solution. Cells were cultured at 37 °C, 5% CO2. The differentiation of IMR-32 cells into neuronal cells was carried out by using 10 µM retinoic acid in 2% FBS for 9 days (42). For phosphoproteomic analysis, the cells were treated with: 1) 100 nM rotenone (PD model) for time points 2,5,15,30 minutes, 2) 100 nM rotenone with 200 µg/ml of *G. glabra*co-treatment for the time points 2, 5, 15, 30 minutes. 3) 200 µg/ml G. glabra extract alone was treated for 30 minutes, and 4) The untreated cells were taken as a control. Cell culture treatments were carried out as biological replicates and processed separately for phosphoproteomic sample preparation.

### Sample preparation for total and phosphoproteomic analysis

The treated cells were washed in ice-cold 1X Phosphate Buffered Saline (PBS) four times, and cells were scraped from the culture dish after the addition of lysis buffer [8 M urea in 200 mM N-(2-Hydroxy ethyl)-Piperazine ethane Sulfonic acid (HEPES) buffer, with sodium pyrophosphate (2.5 mM), sodium orthovanadate (1 mM), and β-glycerophosphate (1 mM)]. The lysate was sonicated using a Q-Sonica probe sonicator at an amplitude of 20% for two cycles, followed by centrifugation at 12,000 × g for 20 minutes at room temperature. The supernatant was transferred to a fresh tube.

Protein concentration was estimated using Bicinchoninic Acid (BCA) assay using Bovine Serum Albumin (BSA) as a standard, and the same was confirmed visually by resolving the samples using Sodium Dodecyl Sulfate-Polyacrylamide Gel Electrophoresis (SDS-PAGE) on a 10% gel. Based on the protein concentrations, 1 mg protein from each condition was taken for further processing. The protein samples were reduced using 10 mM dithiothreitol (DTT) for 1 hour at room temperature and alkylation using 20 mM iodoacetamide (IAA) for 15 minutes, in the dark at room temperature. To reduce the urea concentration, the samples were diluted with 20 mM HEPES to a final urea concentration of 1 M. The samples were digested with Tosyl phenylalanyl chloromethyl ketone (TPCK) treated trypsin, incubated overnight at a ratio of 20:1 (protein: enzyme) at 37 °C. The efficiency of trypsin digestion was evaluated by resolving the pre- and post-digestion samples on 10% SDS-PAGE gels. The peptides were dried overnight in a Speed Vac and stored at -20 °C until further processing.

### Peptide clean-up

To remove urea and salts, the digested peptides were desalted before labeling with tandem mass tags. The peptide samples were acidified with 1% trifluoroacetic acid (TFA). Sep-Pak C-18 cartridge, fitted with a 50 ml syringe, was used for cleanup. The cartridge was activated with 100% acetonitrile (ACN) and equilibrated with 0.1% formic acid (FA). The peptide samples were loaded onto the column. The flow-through was collected and allowed to bind to the column again. It was then washed thrice with 0.1% FA and eluted with 0.1% FA in 40% ACN. The eluate was collected and dried in a Speed Vac and taken for isobaric labeling using Tandem mass tags (TMT).

### Isobaric labeling using TMT-10plex

Peptide samples were resuspended in 50 mM Triethylammonium bicarbonate (TEAB) buffer, and the peptide concentrations were estimated with the Pierce peptide estimation kit. 1 mg of peptide sample from each treatment condition was labeled with TMT 10plex labels. Peptides from the ten different samples were labeled as represented in Table 1. The biological triplicates were independently labeled as per the manufacturer’s protocol. The labels were dissolved in 41 µl of anhydrous acetonitrile and vortexed to ensure dissolution. The corresponding TMT-tags were added to the peptides from the treatment condition for each replicate. The peptide samples with TMT labels were incubated at room temperature for 1h, and the reaction was quenched by adding 8 µl of 5% hydroxylamine. The labeled samples were pooled and dried using Speed Vac. A fraction of the pooled samples was aliquoted for total proteomic analysis. The dried peptide samples were stored in a deep freezer at -20 °C until further phosphopeptide enrichment.

**Table 1.** Details of 10plex TMT labels used for temporal quantitative total and phosphoproteomic analysis.

| Label | Sample |
| --- | --- |
| 126 | Control |
| 127N | 100 nM Rotenone- 2 minutes |
| 127C | 100 nM Rotenone- 5 minutes |
| 128N | 100 nM Rotenone- 15 minutes |
| 128C | 100 nM Rotenone- 30 minutes |
| 129N | 100 nM Rotenone + 200 μg/ml *G. glabra*- 2 minutes |
| 129C | 100 nM Rotenone + 200 μg/ml *G. glabra*- 5 minutes |
| 130N | 100 nM Rotenone + 200 μg/ml *G. glabra*- 15 minutes |
| 130C | 100 nM Rotenone + 200 μg/ml *G. glabra*- 30 minutes |
| 131 | 200 μg/ml *G. glabra* |

### Phosphopeptide enrichment

Phosphopeptide enrichment was carried out with sequential metal oxide affinity chromatography (SMOAC) using immobilized metal oxide (TiO_2_), followed by immobilized metal affinity (Fe-NTA) enrichment, using the manufacturer’s protocol. The TiO_2_ enrichment was carried out using High-Select™ TiO_2_ Phosphopeptide Enrichment Kit (Thermo Fisher Scientific, USA). Briefly, the dried peptide samples were reconstituted in the binding/equilibration buffer and vortexed thoroughly to ensure complete dissolution. The TiO_2_ binding spin column was washed with wash buffer, followed by activation with the binding/equilibration buffer at 3,000 x g for 2 minutes. Upon activation, the pooled TMT-labeled samples were loaded onto the activated columns by centrifugation at 1,000 x g for 5 minutes. The columns were then washed with wash buffer at 3,000 x g for 2 minutes to remove the non-specific binding of peptides, and the flow-through was collected in a fresh tube and stored for further Fe-NTA enrichment. The bound phosphopeptides were eluted using the elution buffer at 1,000 x g for 5 minutes, collected in a new tube, and dried.

The Fe-NTA enrichment was carried out using the High-Select Fe-NTA Phosphopeptide Enrichment Kit (Thermo Fisher Scientific, USA). The Fe-NTA column is shipped with a storage buffer, which was first removed using centrifugation at 1,000 x g for 30 seconds. The column was then equilibrated using a binding/wash buffer. The flow-through samples from the TiO_2_ method were resuspended in binding/wash buffer and loaded onto the column, and incubated for 30 minutes with occasional inverted mixing. The sample with the column was then centrifuged at 1,000 x g for 30 seconds and washed to remove non-specific binding. The phosphopeptides were eluted at 1,000 × g for 30 seconds using the elution buffer, and the procedure was repeated once. The phosphopeptides were collected in a new tube and dried. Prior to fractionation, the phosphor-peptides enriched by both methods were pooled, dried using a SpeedVac, and stored in a -20 °C freezer until further analysis.

### Fractionation of phosphopeptides

The enriched phosphopeptides and peptides for total proteome analysis were reconstituted in 0.2% TFA and fractionated using Sep-Pak C-18 cartridges. The C-18 cartridges were activated with 100 % ACN, followed by equilibration with 0.2% TFA. The peptides were loaded onto the C-18 cartridges, and the flow-through was collected, which was reloaded onto the column to allow binding again. The column was washed with 0.2% TFA and proceeded for fractionation. The peptides were eluted with 0.2% TEAB buffer and different percentages of ACN, varying from 1% to 80%. A total of 24 fractions were prepared, and they were concatenated to a final set of 12 fractions from each of the three biological replicates. The fractions were dried in Speed Vac and stored at -20 °C until further LC-MS/MS analysis.

### Quantitative total proteomic and phosphoproteomic analysis using the LC-MS/MS

LC-MS/MS analysis was carried out on an Orbitrap Fusion Tribrid mass spectrometer (Thermo Fisher Scientific, Bremen, Germany) coupled with Easy-nLC 1200 (Thermo Scientific, Odense, Denmark). The fractionated and dried peptides were reconstituted using 0.1% FA and introduced into the nanoViper trap column (75 µm x 2 cm and 3 µm, C18) (Thermo Fisher Scientific). EASY-Spray C18 PepMep Column (75 µm x 50 cm, 275 µm, 100 Å) maintained at 40 °C was used for peptide resolution. The peptide resolution was carried out using a gradient of 5-35% solvent B (0.1% FA in 80% ACN) at a 300 nl/minute flow rate for 110 minutes. A total run time of 140 minutes was used for the LC-MS/MS analysis, including the column conditioning and sample loading.

The data-dependent acquisition was carried out using the Orbitrap mass analyzer at a resolution of 120,000 at 200 m/z, within a mass range of 400-1,600 m/z. MS/MS fragmentation was done for the most intense precursor ions, at top speed data-dependent mode with a 3 seconds cycle time. Fragmentation was carried out using higher-energy collision dissociation (HCD) mode with a 35% normalized collision energy. HCD was done in the Orbitrap mass analyzer with a scan range of 400-1,600 m/z and a resolution of 60,000 at 200 m/z. The peptide charge states of 2-6 and a dynamic exclusion of 30 seconds were selected with a mass window of 10 ppm. Data acquisition was carried out as biological triplicates.

### 3.9.8 Database search for phosphoproteomic analysis

The mass spectrometry raw data files were processed using Proteome Discoverer, version 2.2 (Thermo Fisher Scientific, Bremen, Germany). The data were searched against the Human protein database RefSeq release 94 and known contaminants (containing 81,096 entries and 116 contaminants)using the SequestHT and MASCOT search algorithms. The minimum peptide length of 7 amino acids, with trypsin as the proteolytic enzyme and one missed cleavage, was set. Precursor and fragment level mass tolerances were selected at 10 ppm and 0.05 Da, respectively. Fixed modifications were as follows: carbamidomethylation of cysteine residues and TMT modification at the peptide N-terminus and lysine residues. Dynamic modifications were protein N-terminal acetylation, phosphorylation at serine (S), threonine (T), and tyrosine (Y), and oxidation of methionine. The ptm-RS node was used for the identification of phosphosites on the S, T, and Y positions, with a phosphorylation cut-off of 75%. The false discovery rate (FDR) was applied at 1% at the peptide and peptide spectrum match (PSM) level at the percolator node, and the data normalization was carried out on the total peptide amount.

### Analysis of phosphoproteomic data

The protein and peptide group output file from the total proteomic and phosphoproteomic experiment in Proteome Discoverer 2.2 was processed in Perseus (27348712, 30931570, 26725330, 23176165), using peptide intensity as the input file. The filter option in Perseus was used to select peptides identified in at least two replicates, which were then considered for analysis. The peptide intensity data were log10-transformed, and the missing values were imputed using the normal distribution curve with the default parameters of 0.3 widths and 1.8 downshift. After replacing the missing values, the data were transformed to their antilog scale, and the fold change (FC) ratios were calculated for rotenone-treated at different time points with respect to control and rotenone +*G. glabra*co-treatment with respect to rotenone at different time points. The FC ratios were log-transformed (log2FC), and the student’s t-test was used for computing the *p-*values. A *p*-value <0.05 was considered significant, and an FC cut-off of ≥1.25 and ≤0.8 was considered up- and downregulated proteins, respectively.

The significantly altered phosphoproteins were subjected to further downstream analysis. Gene ontology (GO) and KEGG pathway analysis for the differentially expressed phosphopeptides and their corresponding proteins were performed using DAVID and Reactome(43–45). To identify the kinases regulating the differentially regulated phosphoproteins and their interactions, Kinase Enrichment Analysis (KEA) was performed using ShinyGo 0.85.1 (46). The kinase-target regulatory network was visualized using Cytoscape 3.8.0 (47,48).

### Data records

The mass spectrometry raw data and the Proteome Discoverer-searched data were submitted to the ProteomeXchange Consortium (http://proteomecentral.proteomexchange.org) via the PRIDE repository (49,50), with the dataset identifier PXD079270.

## Results

### Temporal phosphoproteomic profiling identifies early phosphorylation dynamics regulated by *G. glabra*

To investigate the early phosphorylation-mediated signaling events associated with *G. glabra* -mediated neuroprotection, we performed a temporal quantitative phosphoproteomic analysis in rotenone-treated IMR-32 neuronal cells at 2, 5, 15, and 30 minutes following treatment. A schematic overview of the experimental workflow is illustrated in Figure 1A. Briefly, differentiated IMR-32 cells were treated with rotenone alone or in combination with *G. glabra*extract, followed by TMT-based quantitative total proteomic and phosphoproteomic analysis using LC-MS/MS. Phosphopeptide enrichment was carried out using sequential TiO2 and Fe-NTA enrichment strategies prior to mass spectrometric analysis.

**Figure 1:**
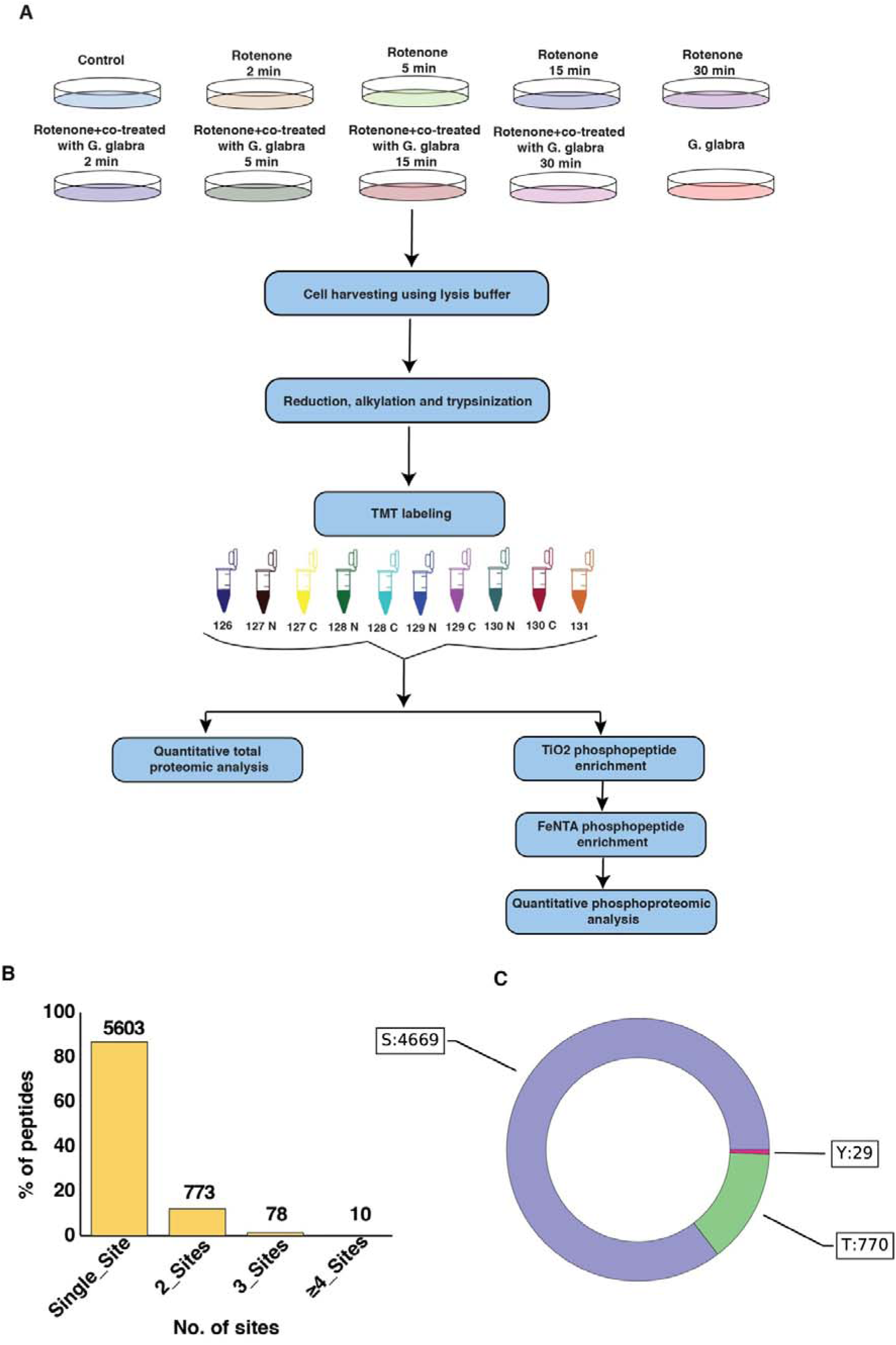
Overview of the phosphoproteomic analysis. **(A) The workflow and experimental design applied for** LC-MS/MS phosphoproteomic analysis. (B) Bar graph representing the number of identified phosphopeptides and their associated phosphorylation sites. (C) Doughnut chart depicting the proportion of phosphopeptides phosphorylated on serine (S), threonine (T), and tyrosine (Y) residues.

The phosphoproteomic analysis identified 6,424 phosphopeptides corresponding to 2,368 phosphoproteins, resulting in the identification of 5,468 phosphorylation sites. After stringent filtering, 4,178 phosphopeptides corresponding to 1,735 phosphoproteins were retained for further analysis. The majority of identified phosphopeptides contained single phosphorylation sites, whereas a smaller proportion represented multiply phosphorylated peptides and were phosphorylated predominantly at serine residues, followed by threonine and tyrosine. (Figure 1B&C). Detailed information on the total number of proteins and phosphopeptides is provided in Supplementary Table 1-2.

### *G. glabra* regulates phosphorylation events disrupted in the rotenone-induced cellular model of Parkinson’s disease

Time-dependent differentialphosphorylation analysis using a FC cutoff of ≥1.25 (upregulated) or ≤0.8 (downregulated) with p < 0.05 revealed extensive temporal phospho-regulation following rotenone exposure and *G. glabra* co-treatment. The time dependent regulation of phosphopeptides and their corresponding phosphoproteins in the table 2. A total of 501 peptides, corresponding to 361 proteins, were dysregulated by rotenone across all the four-time points. Treatment with *G. glabra* showed the restoration of 62 peptides corresponding to 55 proteins and highlighted the additional regulation of 91 peptides corresponding to 75 proteins across the four-time points (Table 2). In summary, *G. glabra*treatment was found to regulate 154 peptides corresponding to 130 proteins across the four-time points.Rotenone treatment induced rapid phosphorylation changes across all early time points, with the highest number of dysregulated phosphopeptides observed at 2 and 5 minutes. In contrast, *G. glabra* co-treatment restored a subset of rotenone-dysregulated phosphorylation events, particularly during the early signaling phase at 2 and 5 minutes **(Table 2)**. The list of differential regulation of phosphoproteins by Rotenone, *G. glabra* co-treatment and *G. glabra* treatment was given in **Supplementary Table 3-5.** Venn diagram analysis demonstrated distinct and overlapping phosphoprotein regulation across the temporal conditions, indicating dynamic signaling rewiring during neuroprotective response **(Figure 2A-D).**The volcano plots showing the differential regulation of the phosphoproteins across the four time points is depicted in **Figure 3A**. A heatmap representing the differential regulation of 150 phosphorylated peptides is summarized in **Figure 3B**.

**Figure 2.**
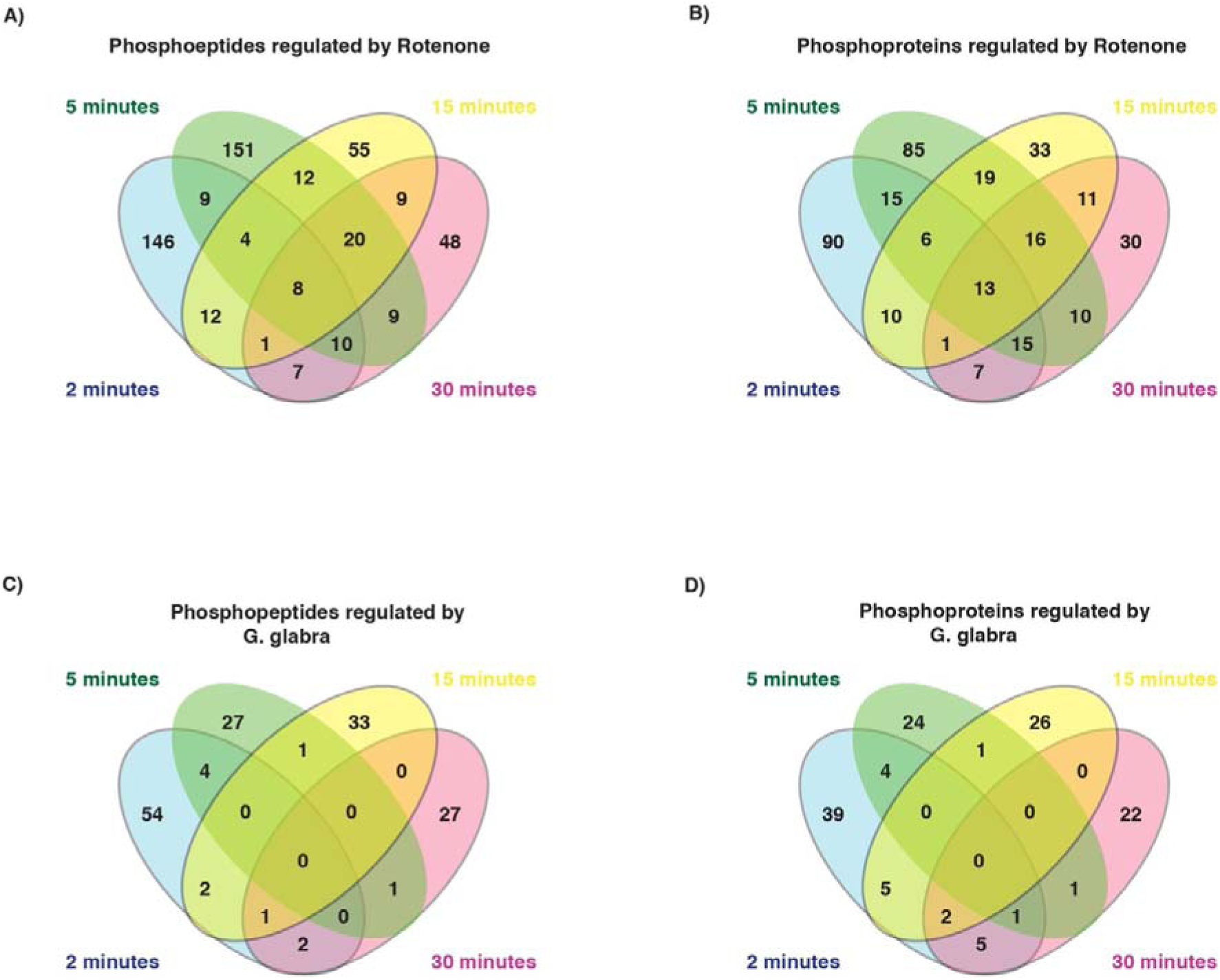
A-D: Venn diagrams showing the temporal comparison of regulated phosphopeptides and proteins across time points under different treatment conditions. (A) phosphopeptides regulated by rotenone, (B) phosphoproteins regulated by rotenone, (C) peptides regulated by combined rotenone and *G. glabra* extract treatment, and (D) proteins regulated by combined rotenone and *G. glabra* extract treatment.

**Figure 3.**
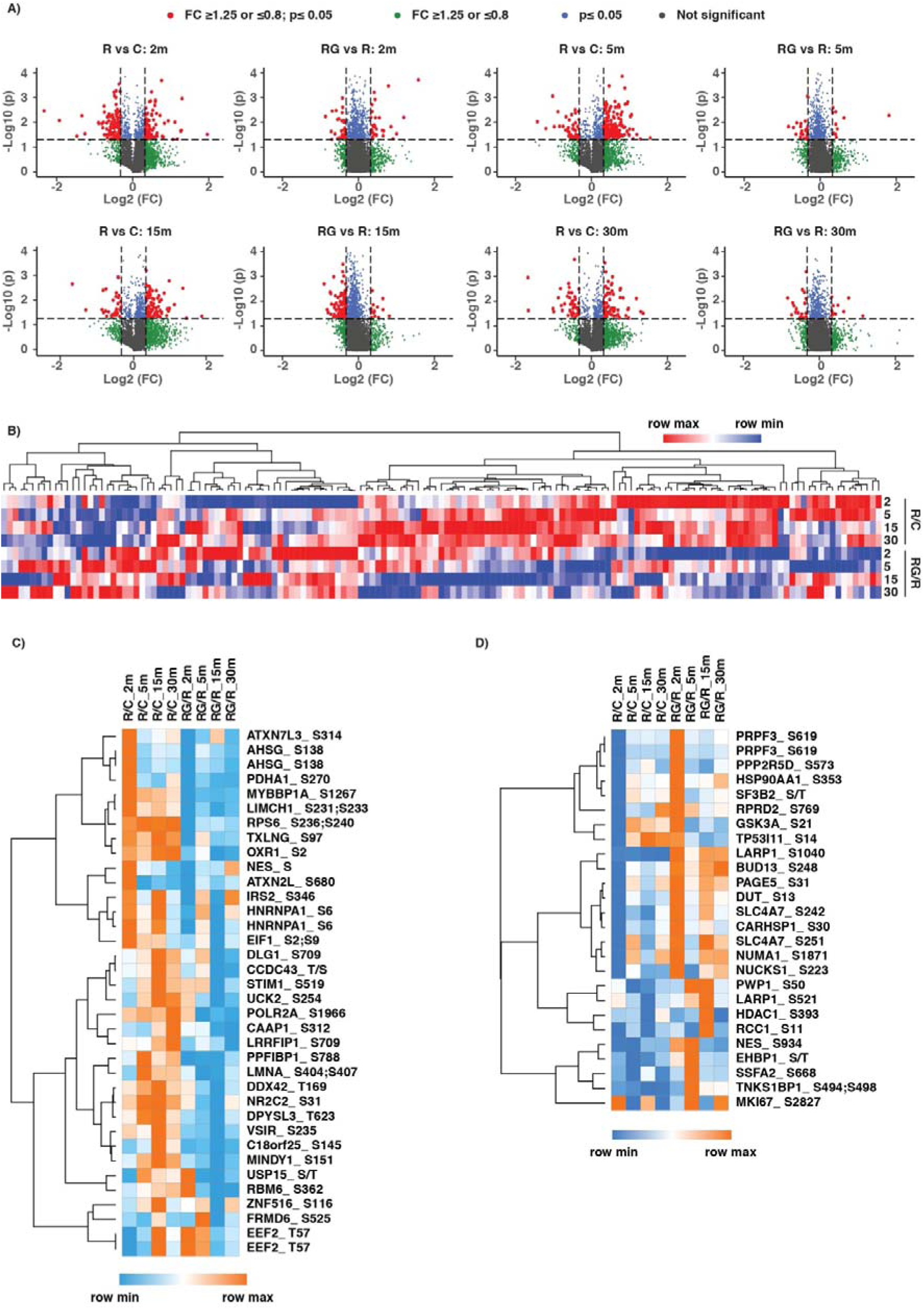
A-D:Differential phosphoprotein expression across treatment groups. (A) Volcano plots depicting differential phosphoprotein abundance for each pairwise comparison at 2, 5, 15, and 30 minutes post-treatment.(B) Heatmap showing normalized phosphopeptide abundance at 2, 5, 15 and 30 min timepoints.(C) Phosphoproteins exhibiting increased phosphorylation upon rotenone treatment and are restored by *G. glabra* treatment. (D) Phosphoproteins exhibiting decreased phosphorylation upon rotenone treatment and are restored by *G. glabra* treatment.

**Table 2:** Temporal regulation of phosphorylation by rotenone and *G. glabra*.

| Time,<br>minutes | Dysregulated by<br>Rotenone |  | Restored by *G. glabra* |  | Additional<br>regulation by *G. glabra* |  |
| --- | --- | --- | --- | --- | --- | --- |
|  | Peptides | Proteins | Peptides | Proteins | Peptides | Proteins |
| 2 | 197 | 157 | 31 | 26 | 33 | 33 |
| 5 | 223 | 179 | 7 | 7 | 27 | 26 |
| 15 | 121 | 109 | 25 | 23 | 13 | 16 |
| 30 | 112 | 103 | 4 | 4 | 32 | 31 |

We have further checked the differentially regulated phosphoproteins along with their phosphosites across the four time points. The heatmap demonstrated that *G. glabra* treatment restored several phosphorylation events altered by rotenone exposure **(Figure 3C-D).** Notably, the majority of restored phosphoproteins were observed during the earliest treatment intervals, suggesting that *G. glabra* exerts rapid effects on kinase-mediated signaling pathways associated with neuronal stress response and survival.

### *G. glabra* restores rotenone-induced phosphorylation alterations during early signaling events

To evaluate the effects of *G. glabra* on early rotenone-induced phosphorylation changes, comparative temporal phosphoproteomic analysis was performed across 2-, 5-, 15-, and 30-minute treatment intervals. Differential phosphorylation analysis revealed that rotenone treatment induced rapid and dynamic alterations in multiple phosphoproteins associated with neuronal stress signaling, cytoskeletal organization, RNA processing, and survival pathways. Interestingly, co-treatment with *G. glabra* restored a substantial proportion of these dysregulated phosphorylation events, particularly during the early signaling phases at 2 and 5 minutes, indicating rapid modulation of kinase-driven cellular responses.

Quantitative phosphoproteomic analysis identified distinct subsets of phosphoproteins that were either upregulated or downregulated following rotenone exposure and subsequently normalized upon *G. glabra* treatment **(Figure 3A-B)**. Several phosphoproteins involved in stress adaptation, cytoskeletal remodeling, RNA metabolism, and kinase signaling demonstrated significant restoration trends across the temporal conditions. Notably, proteins associated with RNA splicing and mRNA processing, including *PRPF3*, *DDX42*, *HNRNPA1*, *SF3B2*, and *POLR2A*, exhibited dynamic phosphorylation changes during the early treatment intervals, suggesting rapid modulation of post-transcriptional regulatory mechanisms.

In addition to spliceosome-associated proteins, several phosphoproteins involved in neuronal survival and signaling pathways showed altered phosphorylation patterns in response to *G. glabra* treatment. These included *GSK3A*, *IRS2*, *HSP90AA1*, AKT-associated signaling proteins, RPS6 family proteins, and *EEF2*, which are known regulators of stress response, protein synthesis, and neuronal survival pathways. Temporal clustering analysis further demonstrated that *G. glabra* -mediated restoration of phosphorylation signatures occurred predominantly during the earliest treatment intervals, supporting the hypothesis that early kinase signaling events play a critical role in the neuroprotective effects of *G. glabra* **(Figure 3B-C)**.

Collectively, these findings suggest that *G. glabra* rapidly counteracts rotenone-induced signaling perturbations through restoration of phosphorylation-mediated cellular pathways associated with neuronal stress adaptation and survival.

### Integrated phosphoproteomic analysis reveals regulation of RNA processing, AMPK signaling, and cell-cycle pathways

To further investigate the biological significance of the phosphorylation changes regulated by *G. glabra* , differentially expressed phosphoproteins were subjected to Gene Ontology (GO), pathway enrichment, and protein interaction network analyses. Functional enrichment analysis revealed significant enrichment of pathways associated with RNA splicing, mRNA processing, chromatin remodeling, transcriptional regulation, and cytoskeletal organization in rotenone treatment and *G. glabra* co-treatment groups**(Figure 4A-B).**The **Supplementary Table 6-9** summarizes the GO and pathway analysis for the proteins altered upon Rotenone treatment and *G. glabra* co-treatment. Among the most significantly enriched biological processes were mRNA splicing via the spliceosome, regulation of alternative mRNA splicing, chromatin organization, and positive regulation of transcription by RNA polymerase II, suggesting extensive modulation of nuclear and post-transcriptional regulatory processes during early neuroprotective signaling.

**Figure 4.**
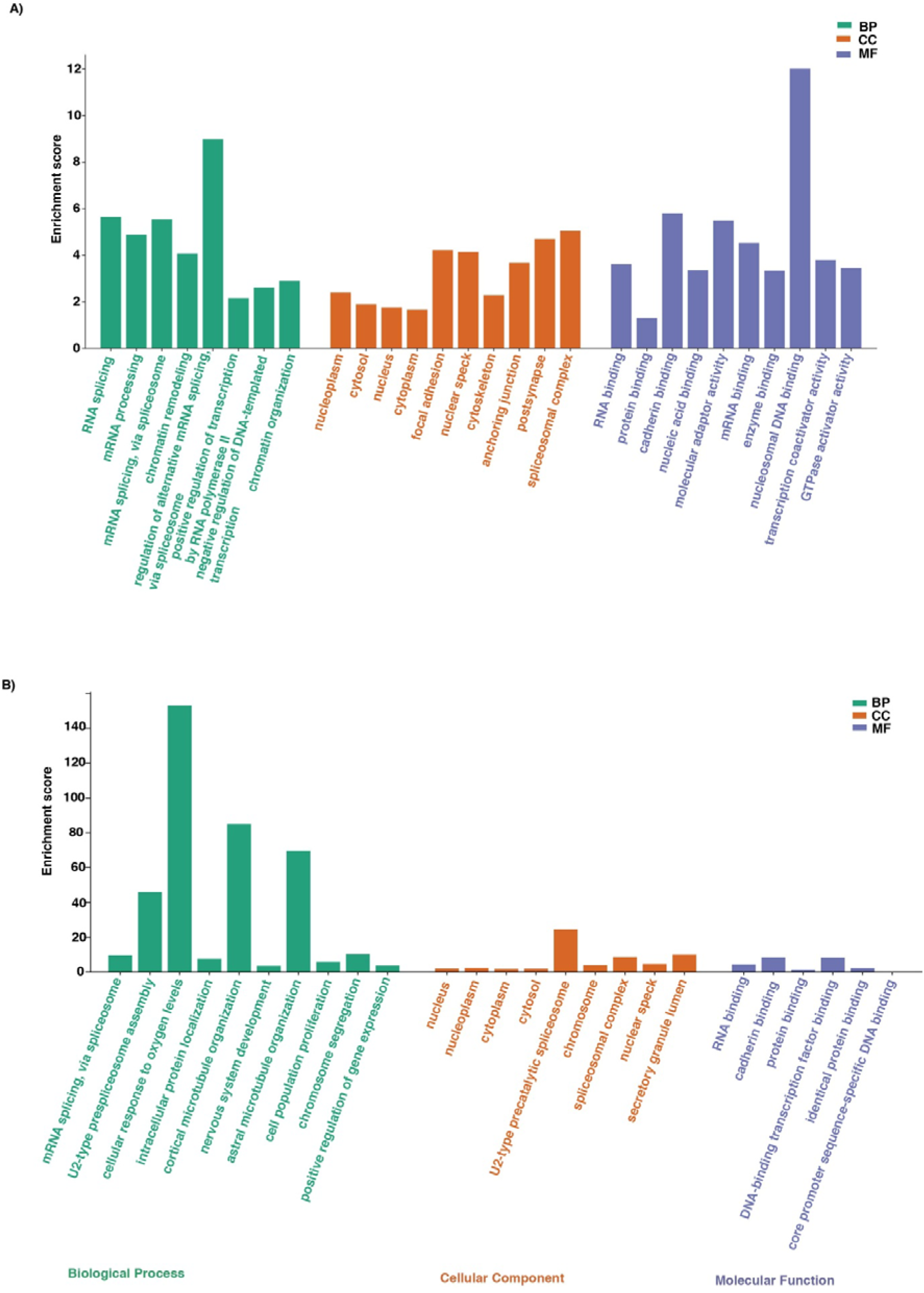
A-B: Gene Ontology (GO) enrichment analysis of proteins altered by Rotenone treatment and *G. glabra* co-treatment. (A) Gene Ontology enrichment analysis of proteins significantly altered upon Rotenone treatment and B) Gene Ontology enrichment analysis of proteins significantly altered upon *G. glabra* co-treatment in Rotenone-treated cells, across three ontology domains: Biological Process (BP), Molecular Function (MF), and Cellular Component (CC).

Cellular component analysis demonstrated strong enrichment of phosphoproteins localized to the nucleus, nucleoplasm, spliceosomal complex, nuclear speckles, and cytoskeleton, while molecular function analysis identified enrichment of RNA binding, cadherin binding, nucleic acid binding, transcription coactivator activity, and molecular adaptor activity. These findings collectively indicate that *G. glabra*-mediated phosphoregulation influences multiple interconnected cellular processes involved in RNA metabolism, intracellular signaling, and structural organization.

Pathway interaction analysis further revealed coordinated regulation of spliceosome-associated pathways, including mRNA splicing, mRNA 3′-end processing, mRNA polyadenylation, and processing of capped intron-containing pre-mRNA **(Figure 5).** Several key phosphoproteins, including *PRPF3, SF3B2, DDX42, HNRNPA1, BUD13,* and *POLR2A*, were repeatedly identified across these pathways, highlighting RNA-processing machinery as a major target of early phosphorylation rewiring following *G. glabra* treatment. These observations suggest that restoration of RNA splicing and transcription-associated pathways may contribute to neuronal adaptation during rotenone-induced stress.

**Figure 5:**
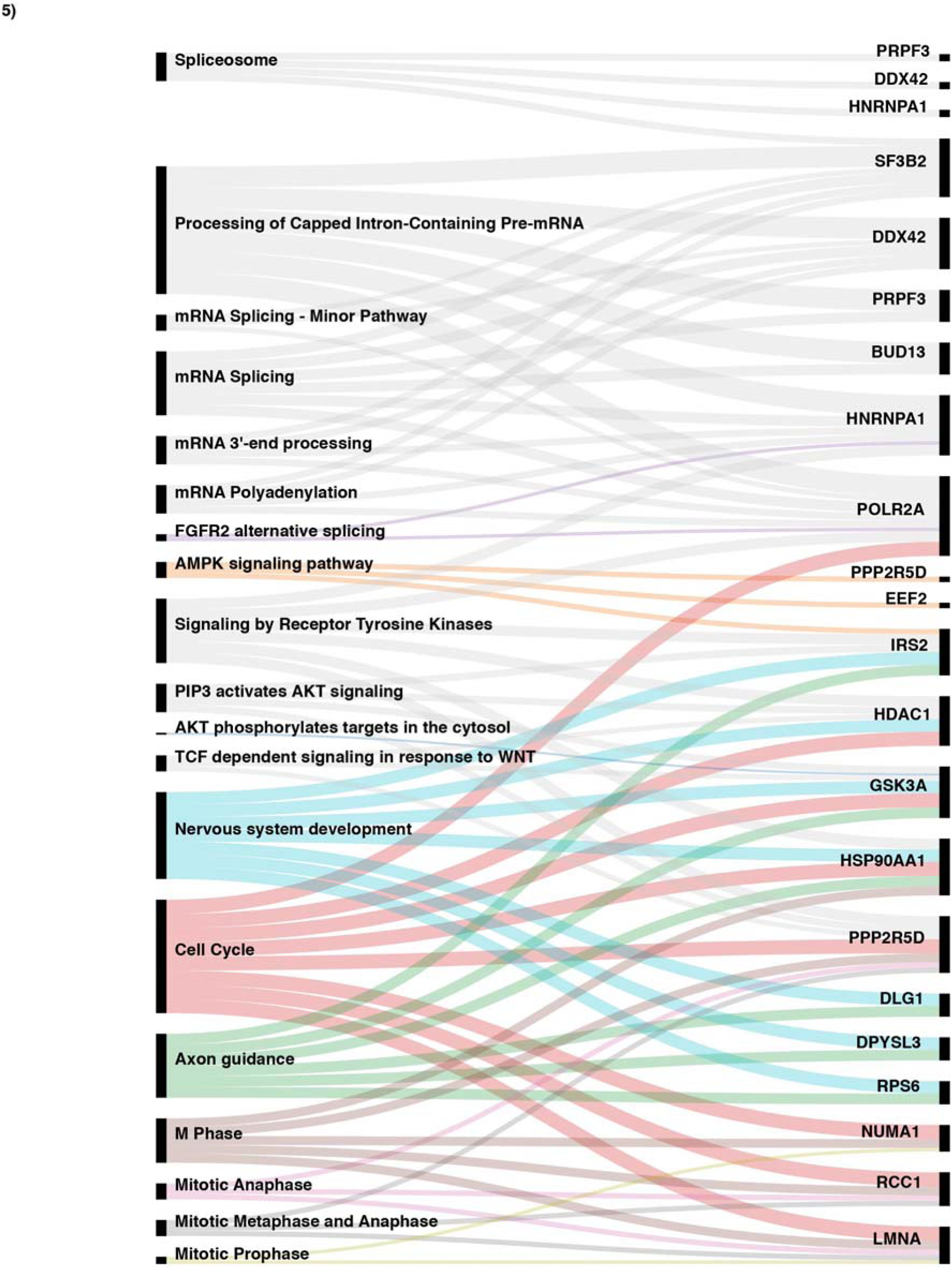
Sankey diagram illustrating shared and distinct pathway enrichment between rotenone-treated and Yashtimadhu co-treated groups.

In addition to RNA-processing pathways, phosphoproteomic analysis identified significant enrichment of AMPK signaling, receptor tyrosine kinase signaling, PIP3-AKT signaling, FOXO-mediated pathways, and cell-cycle associated signaling networks **(Figure 5)**. Multiple phosphoproteins involved in stress adaptation and neuronal survival, including *IRS2,GSK3A, EEF2, HSP90AA1, NUMA1, RCC1, LMNA, and HDAC1*, demonstrated altered phosphorylation patterns following *G. glabra* treatment. Proteins associated with mitotic progression and chromosomal organization such as CDK1, WEE1, and RCC1 were also significantly represented, indicating modulation of cell-cycle-associated stress signaling pathways.

Collectively, these findings demonstrate that *G. glabra*-mediated neuroprotection involves coordinated phosphoregulation of RNA-processing machinery together with restoration of AMPK-, AKT-, and cell-cycle-associated signaling pathways disrupted during rotenone-induced neuronal toxicity.

### Kinase enrichment analysis identifies restoration of AKT/mTOR-associated signaling networks

To identify the upstream kinases associated with the temporal phosphorylation events regulated by Yashtimadhu, kinase enrichment analysis was performed using significantly altered phosphoproteins identified across all treatment conditions and time points. The analysis predicted multiple kinase families associated with rotenone-induced phosphorylation changes and their restoration following *G. glabra* co-treatment. The identified kinases were classified using the human kinome tree, revealing enrichment of several major kinase groups including *AGC, CMGC, STE, CK1, CAMK*, and tyrosine kinase families **(Figure 6).**

**Figure 6:**
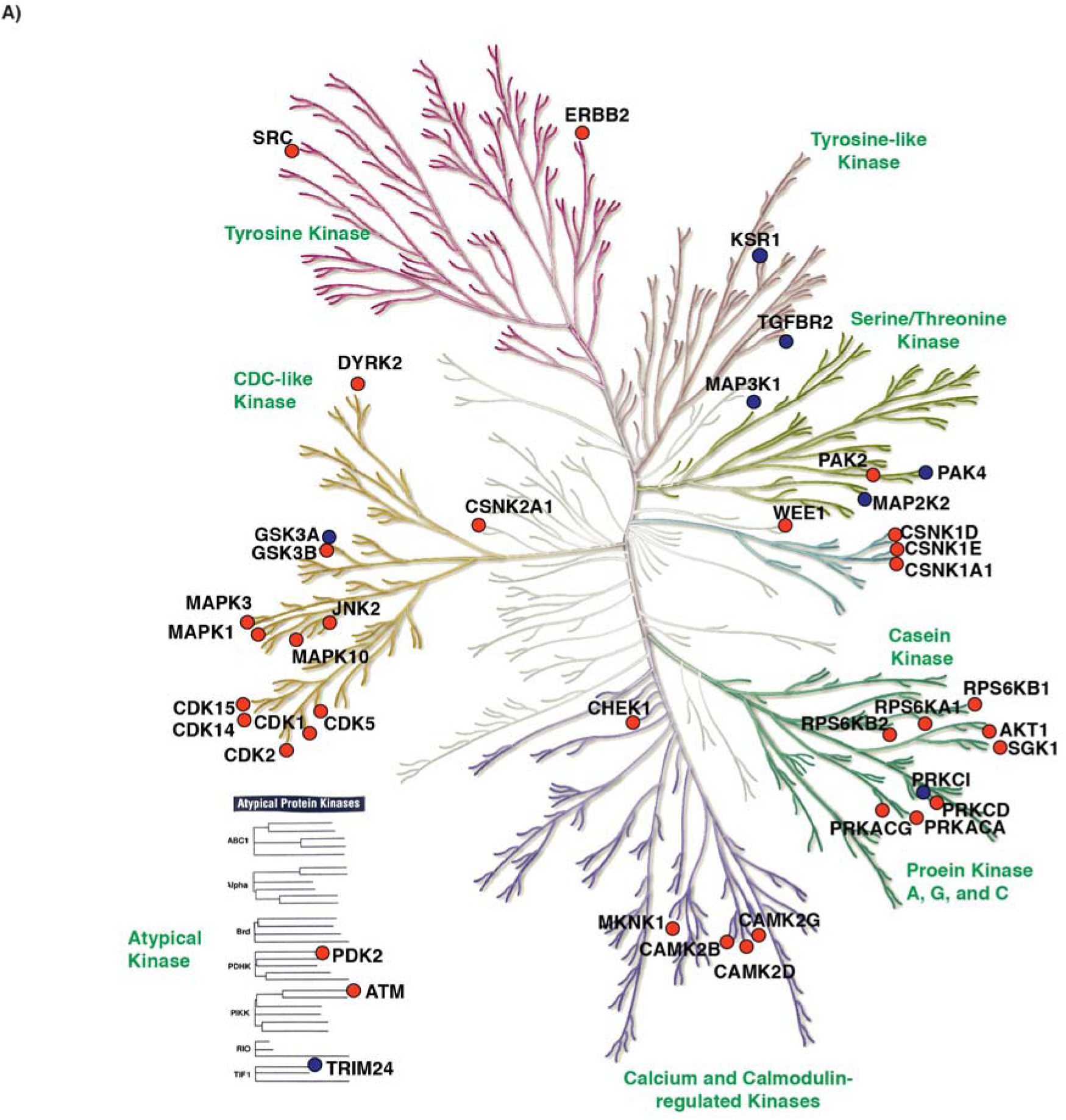
Classification of kinases and kinases identiffied in the dataset represented using human kinome tree. The kinases differentially regulated by *G. glabra* extract co-treatment with rotenone are highlighted in blue and, the enriched upstream kinases are highlighted in red.

Among the enriched kinase families, AGC kinases, including *AKT1, PRKACA, PRKCD, PRKCA, and RPS6KB1/2*, represented one of the most prominent groups associated with *G. glabra*-mediated signaling restoration. In parallel, multiple CMGC family kinases,including*CDK1, CDK2, CDK5, MAPK1, MAPK3, MAPK10, GSK3A, GSK3B, andDYRK2*, were also significantly enriched, indicating coordinated regulation of stress response, neuronal signaling, and cell-cycle-associated pathways. Additional kinase groups, including CAMK family members *CAMK2B, CAMK2D, and CAMK2G*, as well as checkpoint and stress-associated kinases, including *ATM, CHEK1, WEE1*, and *MAPK14*, were identified across the temporal phosphoproteomic datasets. A summary of predicted kinases and their corresponding substrates for rotenone-treated and *G. glabra* co-treatment groups at different time pointsare given in **SupplementaryTable 10-11.**

Kinase-substrate prediction analysis further revealed dynamic regulation of several kinase-associated phosphoproteins involved in neuronal survival, protein translation, stress adaptation, and cytoskeletal organization. Multiple phosphoproteins associated with AKT/mTOR signaling, including *IRS2, RPS6, EEF2, LARP1, and HSP90AA1*, demonstrated altered phosphorylation patterns following rotenone treatment and subsequent restoration upon *G. glabra* co-treatment **(Figure 7A-B).** Furthermore, signaling proteins associated with receptor tyrosine kinase pathways, MAPK signaling, FOXO-mediated pathways, and autophagy-related processes were also enriched, indicating a broad restoration of kinase-mediated cellular signaling networks disrupted during rotenone-induced toxicity.

**Figure 7.**
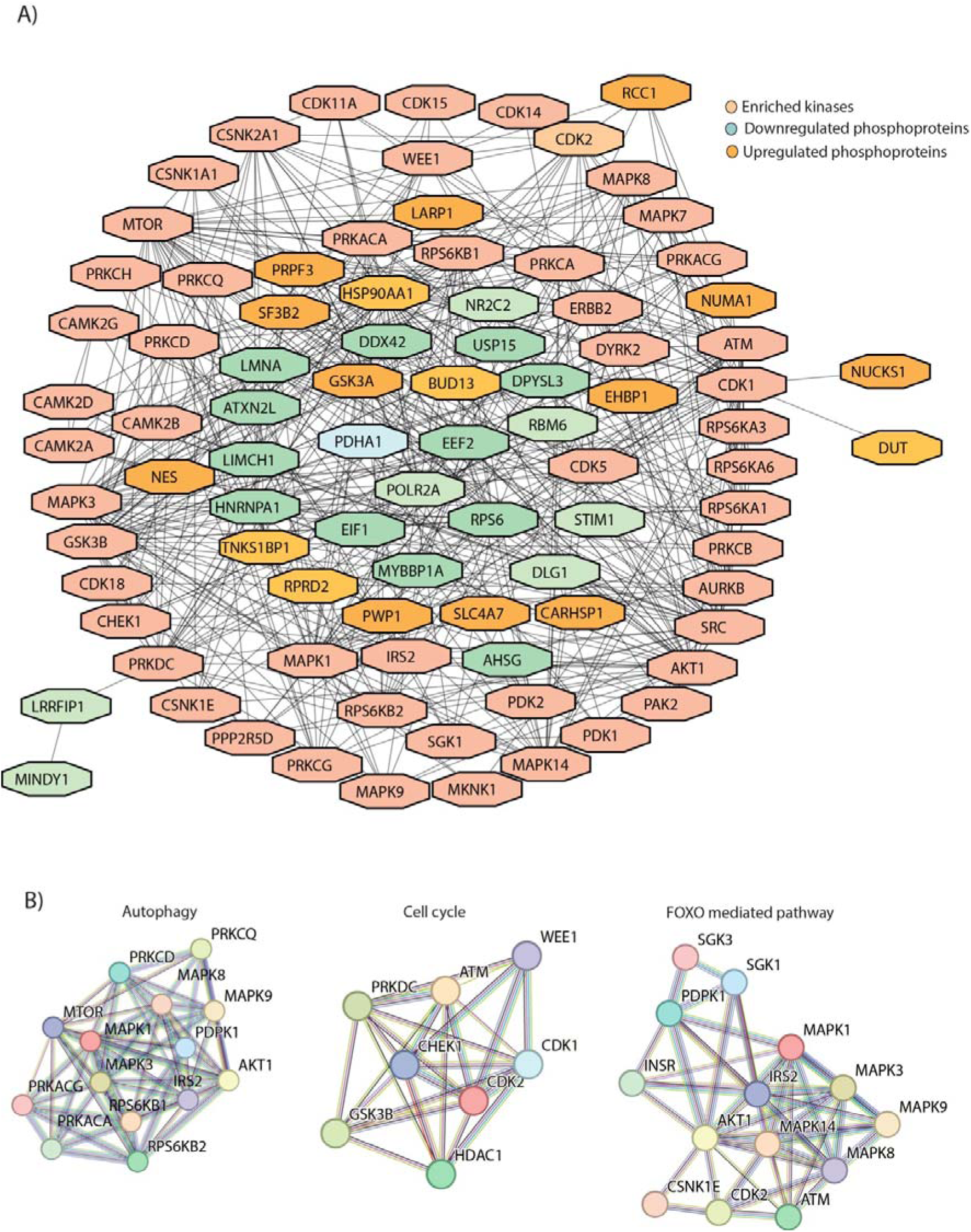
A-B: Protein-protein interaction network of kinases and *G. glabra*-regulated proteins. (A) Kinases and intermediate protein nodes identified in the dataset, and (B) Interactome of the enriched pathways. Node colors indicates the following: Orange- Upregulated phosphoproteins, light blue-Downregulated phosphoproteins, and Red- Kinases.

Interestingly, several kinases associated with neuronal stress adaptation and metabolic regulation, including *AKT1, MTOR, PDPK1, SGK1, MAPK1/3*, and *PRKCD*, were integrated within interconnected signaling networks linked to autophagy, cell-cycle regulation, and FOXO signaling pathways **(Figure 7B).** The enrichment of these kinase networks suggests that *G. glabra* rapidly modulates multiple survival-associated signaling pathways during the early phases of neuronal stress response.

Collectively, these findings demonstrate that *G. glabra*-mediated neuroprotection involves extensive restoration of kinase-centered signaling pathways, particularly those associated with AKT/mTOR signaling, MAPK pathways, stress adaptation, autophagy, and neuronal survival.

### Kinase-substrate interaction analysis reveals coordinated neuroprotective signaling pathways regulated by *G. glabra*

To further elucidate the signaling architecture associated with *G. glabra*-mediated neuroprotection, kinase-substrate interaction networks were constructed using the significantly altered phosphoproteins and their predicted upstream kinases. The interaction analysis revealed extensive connectivity between enriched kinases and phosphoproteins involved in neuronal survival, stress adaptation, autophagy, protein translation, and cytoskeletal organization **(Figure 7A)**. Multiple kinase families including MAPKs, CDKs, AKT-associated kinases, CAMKs, PRKC family members, and checkpoint kinases formed interconnected signaling hubs regulating diverse downstream phosphoproteins across the temporal datasets.The kinase-substrate interaction pathway analysis resulted in the enrichment of160 pathways in KEGG human pathway database with a corrected p-value of ≤0.05 and are provided in **Supplementary Table12.**

Pathway enrichment analysis of the kinase-substrate interaction network identified significant enrichment of autophagy, FOXO signaling, receptor tyrosine kinase signaling, AMPK signaling, insulin signaling, ERBB signaling, and cell-cycle regulatory pathways **(Figure 7B)**. Several key signaling nodes including *AKT1, MTOR, MAPK1/3, PRKCD, PDPK1, SGK1, GSK3A/B, ATM, CDK1, CHEK1*, and *WEE1* were integrated into coordinated signaling modules associated with neuronal stress response and survival. These pathways were connected to phosphoproteins involved in protein synthesis, mitochondrial regulation, synaptic plasticity, and metabolic adaptation.

Notably, proteins associated with translational regulation and metabolic stress response, including *RPS6, EIF4E, EIF4EBP1, EEF2*, and *LARP1,* were linked to mTOR- and AKT-associated signaling modules, suggesting restoration of protein synthesis and cellular energy homeostasis following *G. glabra* treatment. In addition, several phosphoproteins associated with autophagy and stress signaling pathways, including ULK1, PRKAA1, MAPK family proteins, and FOXO-associated regulators, were integrated within the kinase interaction network, indicating modulation of adaptive stress-response pathways disrupted during rotenone exposure.

The interaction network also highlighted extensive crosstalk between receptor tyrosine kinase signaling, MAPK signaling, NF-κB-associated pathways, and cytoskeletal regulatory networks. Proteins including *IRS2, DLG1, JUNB, LMNA, HSP90AA1*, and *CARHSP1* formed central interaction nodes linking neuronal signaling, apoptosis regulation, and cellular stress adaptation. Furthermore, several kinases associated with synaptic plasticity and neuronal differentiation, including CAMK family members and ERBB-associated signaling proteins, were identified within the network, supporting the role of Yashtimadhu in restoration of neuronal signaling homeostasis.

A comprehensive signaling map integrating kinase-substrate interactions across all temporal conditions further demonstrated dynamic pathway rewiring during early neuroprotective response **(Figure 8).** The integrated network revealed coordinated restoration of signaling pathways associated with cellular survival, metabolism, autophagy, RNA processing, and neuronal stress adaptation following *G. glabra* co-treatment. Collectively, these findings indicate that *G. glabra* exerts neuroprotective effects through rapid and coordinated modulation of kinase-driven signaling networks disrupted during rotenone-induced neuronal toxicity.

**Figure 8:**
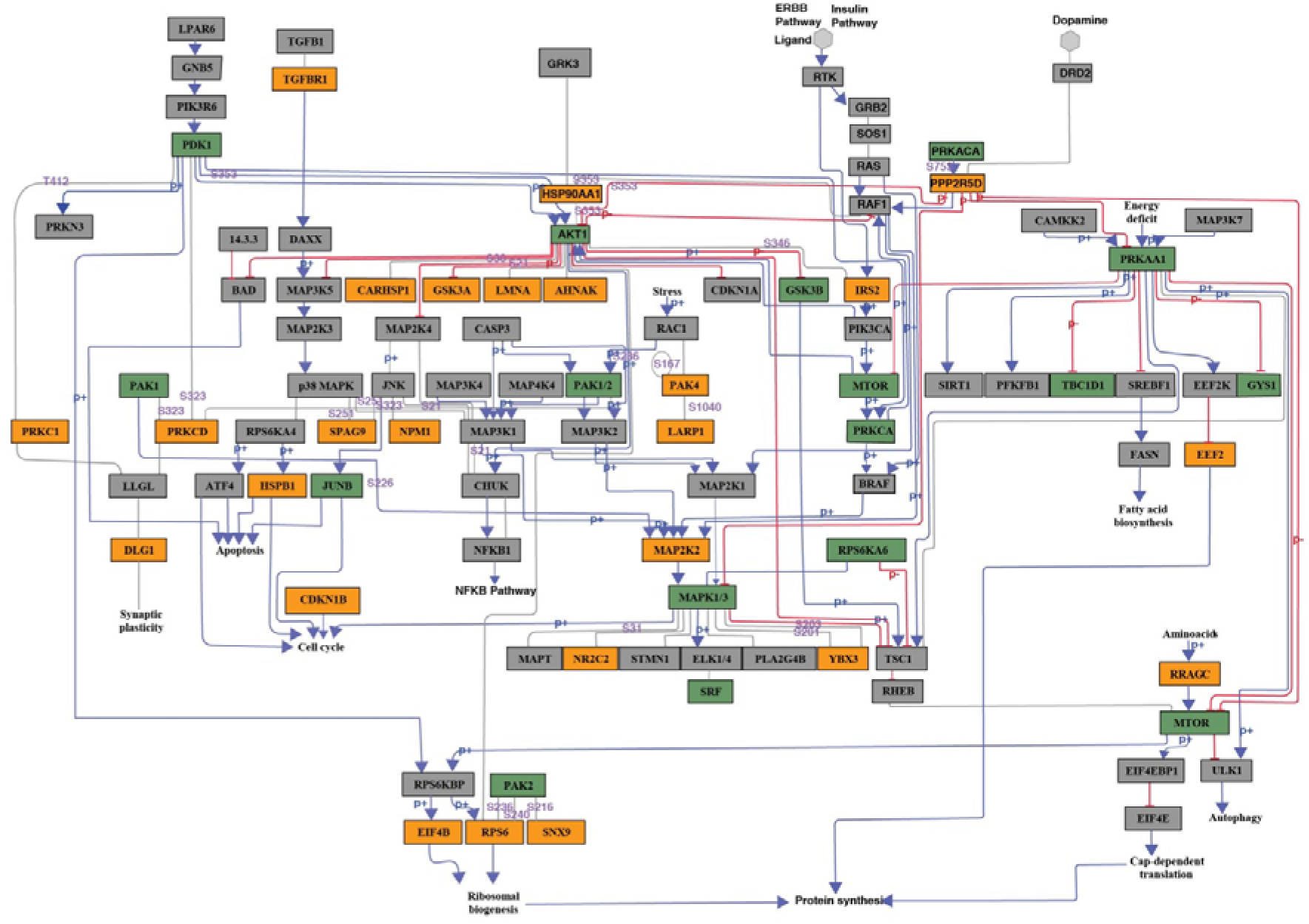
A complete map of the network interaction panning all the time points constructed from the kinase-substrate links and the pathway regulations.

## Discussion and Conclusions

*G. glabra* (Yashtimadhu) has been widely investigated for its neuroprotective, antioxidant, and anti-inflammatory properties in both in vitro and in vivo models of neurodegeneration(14,15). Several studies have demonstrated the therapeutic potential of crude root extracts or bioactive metabolites derived from *G. glabra* in mitigating neuronal stress and improving cellular survival(14,15,18,51,52). In our previous studies, we demonstrated that metabolites present in *G. glabra* powder interact with multiple cellular target proteins, including several kinases and signaling molecules associated with neuronal survival pathways(53). We have also previously reported the global proteomic alterations induced by rotenone exposure and their restoration following *G. glabra* treatment(16). Despite these observations, the early phosphorylation-mediated signaling mechanisms underlying *G. glabra* -induced neuroprotection remained poorly understood. Therefore, in the present study, we employed a temporal quantitative phosphoproteomic approach to systematically characterize the rapid phosphorylation dynamics associated with rotenone-induced neuronal stress and their restoration following *G. glabra*treatment.

Protein phosphorylation is one of the earliest cellular responses following exposure to physiological or pathological stimuli and plays a critical role in regulating signal transduction, protein activity, stress adaptation, and neuronal survival(34,37,38,54–56). Dysregulation of phosphorylation-mediated signaling pathways has been extensively implicated in PD pathogenesis, particularly pathways associated with mitochondrial dysfunction, oxidative stress, autophagy, protein aggregation, and apoptosis(57,58). In the present study, we focused on capturing early temporal phosphorylation events occurring within 2, 5, 15, and 30 minutes following rotenone exposure and *G. glabra* co-treatment in differentiated IMR-32 neuronal cells. Our analysis identified extensive temporal phosphoregulation involving more than 130 phosphoproteins along with dynamic alterations in multiple kinase-associated signaling pathways.

One of the major observations from this study was the rapid restoration of kinase-mediated signaling pathways disrupted during rotenone-induced stress. Kinase enrichment analysis identified several major kinase families associated with *G. glabra*-mediated signaling restoration, including *AGC, CMGC, CAMK, CK1*, and tyrosine kinase families. Among these, *AKT1, MTOR, MAPK1/3, PRKACA, PRKCD, RPS6KB1/2*, and *GSK3A/B* were major signaling nodes restored following *G. glabra* treatment. These kinases are well-known regulators of neuronal survival, protein synthesis, stress adaptation, autophagy, and mitochondrial homeostasis and have been previously implicated in Parkinson’s disease pathogenesis(57–60). Dysregulation of AKT/mTOR and MAPK signaling pathways contributes to neuronal degeneration through impaired energy metabolism, defective autophagy, oxidative stress, and apoptosis. The restoration of these signaling networks observed in the present study suggests that *G. glabra* rapidly modulates multiple pro-survival kinase pathways during the early phases of neuronal stress response.

Interestingly, integrated phosphoproteomic and pathway enrichment analyses also revealed extensive regulation of RNA-processing and spliceosome-associated pathways. Several phosphoproteins involved in RNA splicing and transcriptional regulation, including *PRPF3, SF3B2, DDX42, HNRNPA1, BUD13*, and *POLR2A,* demonstrated dynamic phosphorylation changes across the temporal conditions. Emerging evidence suggests that alterations in RNA metabolism, spliceosome dynamics, and post-transcriptional regulation contribute significantly to neurodegenerative disorders, including Parkinson’s disease. The observed restoration of spliceosome-associated signaling pathways by *G. glabra* may therefore represent an additional neuroprotective mechanism associated with maintenance of transcriptional and translational homeostasis during neuronal stress.

In addition to RNA-processing pathways, several signaling pathways associated with AMPK signaling, FOXO signaling, receptor tyrosine kinase signaling, autophagy, and cell-cycle regulation were enriched in the kinase-substrate interaction network(61–64). Proteins, including *IRS2, EEF2, HSP90AA1, RPS6, LARP1, NUMA1, RCC1, HDAC1*, and *LMNA*, demonstrated altered phosphorylation patterns following rotenone exposure, and subsequent restoration upon *G. glabra* treatment. These signaling pathways are closely associated with cellular energy homeostasis, protein translation, mitochondrial function, and neuronal stress adaptation. Notably, aberrant activation of cell-cycle-associated signaling in post-mitotic neurons has been implicated in neuronal death and neurodegeneration. The modulation of CDK1-, WEE1-, and GSK3-associated signaling observed in the present study suggests that *G. glabra* may attenuate stress-associated cell-cycle dysregulation during rotenone-induced neuronal toxicity.

The kinase-substrate interaction networks generated in this study further highlighted extensive crosstalk between AKT/mTOR signaling, MAPK pathways, autophagy-associated signaling, receptor tyrosine kinase pathways, and stress-response networks. The integration of these phosphorylation events enabled the construction of an early signaling map associated with *G. glabra*-mediated neuroprotection. These findings collectively suggest that *G. glabra* exerts rapid neuroprotective effects through coordinated modulation of phosphorylation-driven signaling pathways that regulate neuronal survival, protein synthesis, autophagy, RNA processing, and metabolic adaptation.

Although the present study provides a comprehensive overview of the early phosphoproteomic alterations associated with *G. glabra*-mediated neuroprotection, certain limitations should be acknowledged. The findings are primarily based on phosphoproteomic and bioinformatic analyses and require further functional validation using targeted biochemical and molecular approaches. Validation of key signaling molecules and phosphorylation events using western blotting, kinase assays, or inhibition studies will further strengthen the mechanistic interpretation of the identified signaling pathways.

In conclusion, this study provides a comprehensive temporal phosphoproteomic landscape of early signaling events associated with *G. glabra*-mediated neuroprotection in a rotenone-induced Parkinson’s disease model. The study highlights the rapid restoration of AKT/mTOR-, MAPK-, AMPK-, and RNA-processing-associated signaling pathways that were disrupted during neuronal stress. The identified kinase-substrate interaction networks and enriched signaling pathways provide important molecular insights into the neuroprotective mechanisms of *G. glabra* and may serve as potential targets for future therapeutic validation in PD.

